# The landscape of incident disease risk for the biomarker GlycA and its mortality stratification in angiography patients

**DOI:** 10.1101/280677

**Authors:** Johannes Kettunen, Scott C. Ritchie, Olga Anufrieva, Leo-Pekka Lyytikäinen, Jussi Hernesniemi, Pekka J. Karhunen, Pekka Kuukasjärvi, Jari Laurikka, Mika Kähönen, Terho Lehtimäki, Aki S. Havulinna, Veikko Salomaa, Satu Männistö, Mika Ala-Korpela, Markus Perola, Michael Inouye, Peter Würtz

## Abstract

Integration of systems-level biomolecular information with electronic health records has led to the discovery of robust blood-based biomarkers predictive of future health and disease. Of recent intense interest is the GlycA biomarker, a complex nuclear magnetic resonance (NMR) spectroscopy signal reflective of acute and chronic inflammation, which predicts long term risk of diverse outcomes including cardiovascular disease, type 2 diabetes, and all-cause mortality. To systematically explore the specificity of the disease burden indicated by GlycA we analysed the risk for 468 common incident hospitalization and mortality outcomes occurring during an 8-year follow-up of 11,861 adults from Finland. Our analyses of GlycA replicated known associations, identified associations with specific cardiovascular disease outcomes, and uncovered new associations with risk of alcoholic liver disease (meta-analysed hazard ratio 2.94 per 1-SD, P=5×10^-6^), chronic renal failure (HR=2.47, P=3×10^-6^), glomerular diseases (HR=1.95, P=1×10^-6^), chronic obstructive pulmonary disease (HR=1.58, P=3×10^-5^), inflammatory polyarthropathies (HR=1.46, P=4×10^-8^), and hypertension (HR=1.21, P=5×10^-5^). We further evaluated GlycA as a biomarker in secondary prevention of 12-year cardiovascular mortality in 900 angiography patients with suspected coronary artery disease. We observed hazard ratios of 4.87 and 5.00 for 12-year mortality in angiography patients in the fourth and fifth quintiles by GlycA levels demonstrating the prognostic potential of GlycA for identification of high mortality-risk individuals. Both GlycA and C-reactive protein had shared as well as independent contributions to mortality hazard, emphasising the importance of chronic inflammation in secondary prevention of cardiovascular disease.

## Introduction

The identification of predictive biomarkers for disease risk is fundamental to precision medicine^1,2^. High-throughput nuclear magnetic resonance (NMR) metabolomics profiling of prospective cohorts have recently revealed that elevated blood concentrations of glycoprotein acetyls (GlycA) are a strong predictive biomarker for long-term risk of morbidity and mortality from diverse diseases, many of which are associated with chronic inflammation^3,4^. GlycA levels have been found to predict risk from cardiovascular diseases^5–8^, certain cancers^6,7,9^, type 2 diabetes^10,11^, non-alcoholic fatty liver^12^, chronic inflammatory conditions^7^, severe infections^13^, and all-cause mortality^6,7^. GlycA itself is a heterogeneous NMR signal quantifying the combined levels of at least five circulating glycoproteins^13–15^ and has long been known to elevate with acute inflammation^14–19^. Recently, in asymptomatic individuals GlycA has been found to denote subclinical chronic inflammation^13^.

Therapeutics targeting inflammation have been of interest for their potential to reduce chronic disease risk^20^. Recently, clinical trials found the anti-inflammatory drug Canakinumab, a monoclonal antibody for IL-1β, significantly reduced incidence and mortality from recurrent cardiovascular events and from lung cancer in patients with previous myocardial infarction and elevated levels of high-sensitivity C-reactive protein (CRP)^21,22^. Chronic inflammation is an important factor in the pathogenesis and severity of numerous age-related conditions and diseases^23,24^ and elevated GlycA has also been found to predict disease severity in patients with rheumatoid arthritis and psoriasis^16–18^, both chronic inflammatory diseases. In psoriasis patients, who are at high risk for cardiovascular disease, GlycA levels were also a predictive biomarker for vascular inflammation and coronary artery disease burden^18^. Importantly, in many studies, GlycA’s predictive effect has been shown to be independent of CRP, the current standard biomarker for inflammation. GlycA represents a promising biomarker to complement CRP in identifying those individuals with chronic inflammation and at increased risk of incident disease or poor prognosis. However, despite the above-mentioned advances, a systematic and unbiased evaluation of the spectrum of cause-specific death and hospitalisation associations for GlycA has been lacking. Such a mapping of disease specificity would clarify the translational and therapeutic applications of GlycA for risk prediction and prognostics.

Here, we systematically evaluate GlycA as a biomarker for future risk of hospitalization or death across the spectrum of all common diseases of two independent population-based cohorts with complete 8-year electronic health record (EHR) follow-up. We further demonstrate the potential of chronic inflammation for stratifying long-term risk of cardiovascular mortality in a cohort of patients with suspected heart attack. Our findings identify several diseases for which GlycA could enable better risk assessment in primary prevention and suggest potential for GlycA in secondary prevention of angiography patients.

## Results

To elucidate the cause-specificity of the disease burden predicted by elevated GlycA, we utilized high-throughput NMR spectroscopy data with matched electronic health records (EHRs) obtained from national care register for health care and causes of death registries from the independent population-based FINRISK97 (N=7,321) and DILGOM (N=4,540) cohorts^25–27^ (**Methods**). Baseline cohort characteristics are reported in **Table 1**. EHRs were collated into distinct outcomes by ICD10 codes at three-digit accuracy and ICD10 disease groups (**Methods**). Only the first incident of each outcome (either a hospital discharge diagnosis or mortality) was considered for each individual. Main and side causes of diagnosis were treated equally when collating outcomes. Participants with prevalent events were excluded for the analysis each outcome. We tested GlycA for association with all 468 disease outcomes that had > 10 incident cases in both DILGOM and FINRISK97 (full listing in **Table S1**) over an 8-year follow-up period (the maximum follow-up time available for DILGOM). Cox proportional hazards models were fit for each outcome separately with age as time scale, and adjusting for smoking, body mass index (BMI), sex, systolic blood pressure, alcohol consumption as well as three all-cause mortality biomarkers previously identified alongside GlycA in FINRISK97: citrate, very-low density lipoprotein (VLDL) particle size, and albumin^6^. Individuals with any previous incidents of the same outcome within the 10 years prior to sample collection for FINRISK97 and 20 years before sample collection for DILGOM (hereby prevalent cases) were excluded when analysing each outcome. We considered GlycA to be biomarker for any outcome which was nominally significant (P<0.05) in both FINRISK97 and DILGOM, and statistically significant after correcting for multiple testing (P<1.1×10^-4^; adjusting for the 468 tested outcomes) in an inverse-variance weighted meta-analysis of FINRISK97 and DILGOM (**Figure 1**).

**Table 1:** Cohort characteristics. ^*^ Not from complete data, systolic blood pressure N=892, current smoking and T2DM N=899. ¶ please see methods for details.

|  | **FINRISK97** | **DILGOM** | **ANGES** |
| --- | --- | --- | --- |
| Number of participants | 7,321 | 4,540 | 900 |
| Number (and %) of women | 3,644 (50%) | 2,387 (53%) | 324 (36%) |
| Mean age in years (and range) | 48 (25–74) | 52 (25–74) | 62 (34-87) |
| Body mass index (kg/m<sup>2</sup>, mean, sd) | 26.6 ± 4.5 | 27.2 ± 4.8 | 28.0 ± 4.3 |
| GlycA (mean, sd) | 1.41 ± 0.25 | 1.30 ± 0.20 | 1.33 ± 0.21 |
| Systolic blood pressure (mean, sd) | 136 ± 20 | 137 ± 20 | 144 ± 22* |
| Alcohol consumption (median, iqr) | 25.4 (3.8-76.6) | 25.1 (5.0-85.2) | NA¶ |
| Current smoking | 1775 (24%) | 835 (18%) | 122 (14%)* |
| Type 2 diabetes | 440 (6%) | 438 (9%) | 238 (26%)* |

**Figure 1.**
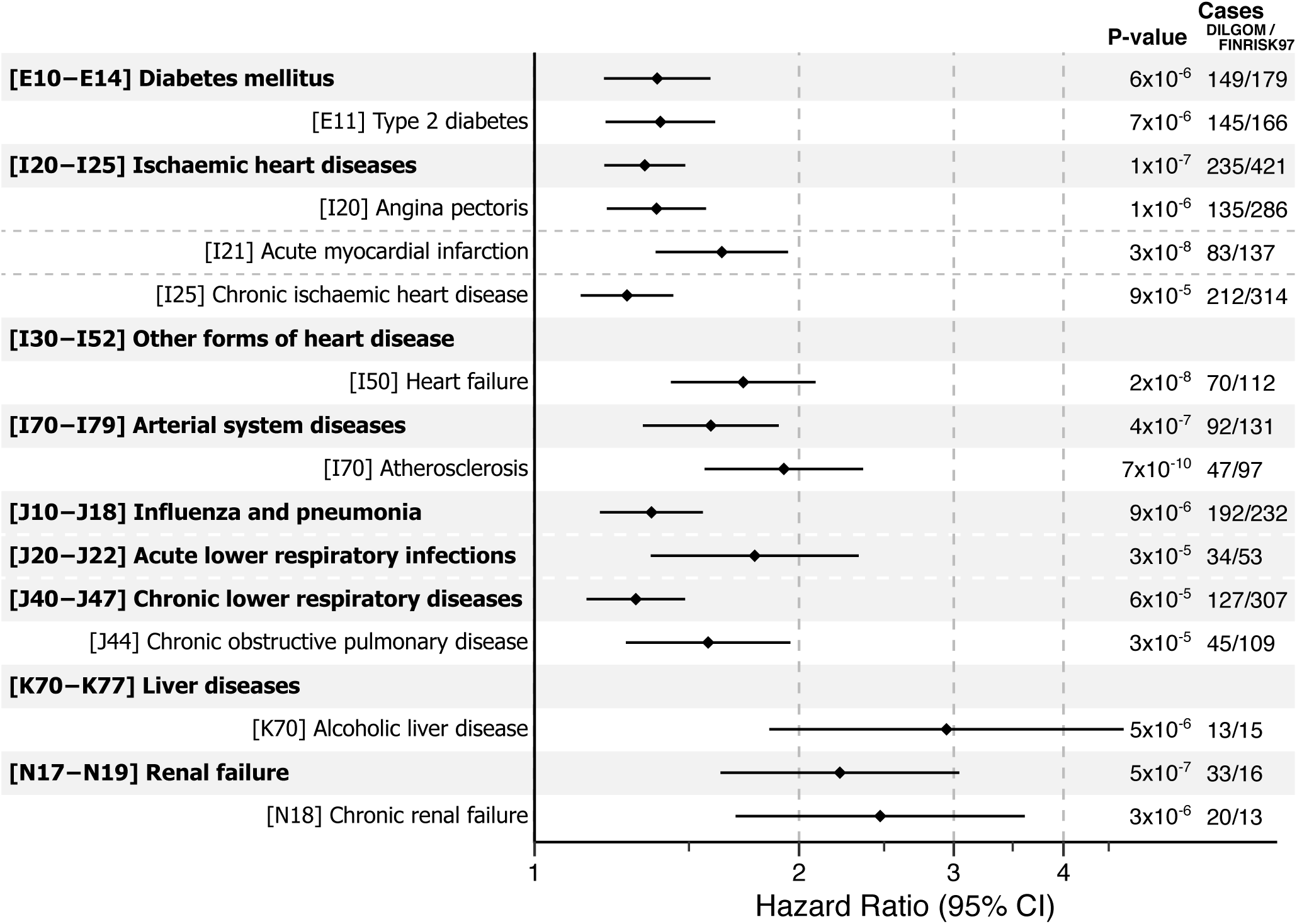
Associations between GlycA and 8-year disease risk assessed by electronic health records.

In total, we observed strong and consistent associations between elevated GlycA concentrations and increased risk for 16 disease outcomes: seven ICD10 categories and nine ICD10 codes at three-digit accuracy (**Figure 1**, **Figure S1**, **Table S2**). Consistent with previous biomarker studies of GlycA, we observed significant and replicable associations with cardiovascular diseases, type 2 diabetes, and respiratory infections (**Figure 1**). Interestingly, the strongest replicable associations between GlycA and 8-year disease risk were with alcoholic liver disease (meta-analysis HR=2.94 per s.d. increase in GlycA, P=5×10^-6^) and chronic renal failures (HR=2.47, P=3×10^-6^), neither of which have yet been investigated by any GlycA biomarker study. Elevated GlycA was also associated with increased 8-year risk of chronic obstructive pulmonary disease (HR=1.58, P=3×10^-5^).

Previous studies of GlycA as a biomarker for cardiovascular disease have used broad definitions encompassing a variety of diseases of the cardiovascular system^5–7^. From our analyses, we were able to identify the specific cardiovascular diseases predicted by GlycA. GlycA was associated with increased risk of atherosclerosis (HR=1.92, P=7×10^-10^), heart failure (HR=1.73, P=2×10^-8^), acute myocardial infarction (HR=1.63, P=3×10^-8^), angina pectoris (HR=1.38, P=1×10^-6^), and chronic ischaemic heart disease (HR=1.27, P=9×10^-5^) (**Figure 1**, **Figure S1**, **Table S2**).

Hazard ratios for GlycA were not attenuated when adjusting models for prevalent cases of each outcome as a covariate rather than excluding them (**Figure S2**). Adjustment rather than exclusion of prevalent cases led to an increase in power for several outcomes, resulting in additional significant associations (**Figure S3**, **Table S3**). We recovered previously observed associations between elevated GlycA and intestinal infections^13^ as well as hypertension^8^, and observed novel associations with 8-year risk of inflammatory polyarthropathies (HR=1.46, P=4×10^-8^), and glomerular diseases (HR=1.95, P=1×10^-6^). Hazard ratios were only weakly attenuated when adjusting for CRP (**Figure S4**), consistent with previous observations that GlycA is largely independent of CRP and may be reflective of a more diverse component of the inflammatory response. The statistical models were stable when comparing the minimally adjusted models (age and sex only) to fully adjusted model (**Figure S5**).

To corroborate the validity of EHR-wide analyses, we compared the disease associations results to adjudicated endpoints for major common diseases (cardiovascular disease subtypes, type 2 diabetes, chronic obstructive pulmonary disease, kidney failure, and any cancer) collated based on health information across national health registries as well as reimbursement information. Our results show consistent associations of EHR mining with those from predefined and widely used adjudicated endpoints (**Methods**, **Figure S6**, **Table S4**).

Finally, we investigated the potential utility of GlycA as a biomarker in secondary prevention in 900 angiography patients from the ANGES study^28^ over a 12-year follow-up period (**Methods**). Baseline cohort characteristics are described in **Table 1**. Their baseline mortality risk over the 12-year follow-up was 27% for all-cause mortality and 14% for cardiovascular mortality. We found GlycA was strongly associated with both 12-year all-cause mortality (HR=1.43, 95% CI: 1.21–1.70, P=4×10^-5^, N=881 with 240 deaths) and cardiovascular mortality risk (HR=1.84, 95% CI: 1.47–2.30, P=8×10^-8^, N=900 with 128 deaths), using follow-up as the time-scale and adjusting for significant covariates (**Methods**). When partitioning the angiography patients by GlycA quintiles we observed a prominent increase in cardiovascular mortality risk when the quintiles were added to the multivariable model (**Figure 2, Table S5**). Strikingly, angiography patients in the fourth and fifth GlycA quintiles were at 4.87 and 5.00 times increased risk of cardiovascular mortality when compared to those in the lowest quintile. In terms of absolute risk there was an increase in 16% between the bottom and top GlycA quintiles for cardiovascular mortality.

**Figure 2.**
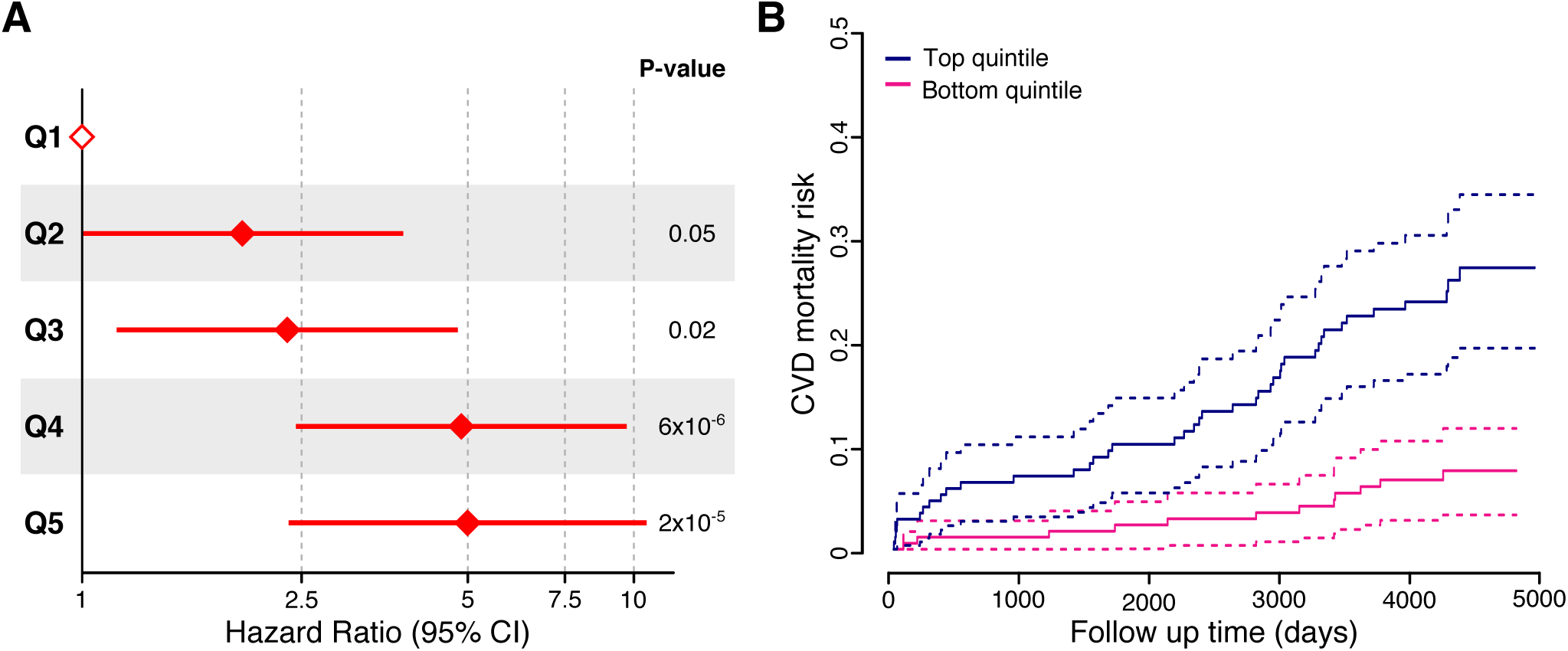
12-year cardiovascular mortality associations by quintiles of GlycA in the ANGES cohort of angiography patients.

We further compared the predictive ability of GlycA and CRP for cardiovascular mortality among patients undergoing angiography. GlycA and CRP were both significantly associated with cardiovascular mortality when analysed separately in covariate-adjusted models (HR=1.69, 95% CI: 1.32–2.16, P=3×10^-5^ and HR=1.61, 95% CI: 1.33–1.96, P=1×10^-6^, respectively). When the two inflammatory markers were included simultaneously in the multivariable model, the hazard ratios for both GlycA and CRP were attenuated but remained strongly predictive (HR=1.37, 95% CI: 1.04–1.81, P=2×10^-2^ and HR=1.45, 95% CI: 1.17–1.80, P=8×10^-4^, respectively). When partitioning the angiography patients by CRP quintiles we observed an increase in cardiovascular mortality risk across increasing CRP concentrations (**Table S6**). Angiography patients in the fourth and fifth CRP quintiles were at 3.12 and 4.61 times increased risk of cardiovascular mortality when compared to those in the lowest quintile.

## Discussion

GlycA is a heterogeneous biomarker associated with both acute and chronic inflammation that has been of intense recent interest as a biomarker for diverse disease outcomes both in asymptomatic people and in those with pre-existing chronic disease^3^. In this study, we performed the first systematic evaluation of GlycA as a reproducible biomarker for onset of all common disease outcomes, using data from 11,861 adults from two independent population-based cohorts with complete EHR over an 8-year follow-up period. In addition to recapitulating previously observed GlycA associations^5–7,10,11,13^, we observed novel associations between elevated GlycA and 8-year risk of alcoholic liver disease, chronic renal failure, glomerular diseases, chronic obstructive pulmonary disease, and inflammatory polyarthropathies, and identified the specific cardiovascular diseases underlying the established GlycA-associated risk of overall cardiovascular disease risk. We further showed potential utility of GlycA for stratifying a 5-fold risk of cardiovascular mortality among 900 angiography patients already at elevated risk for recurrent severe cardiovascular disease events.

We examined the landscape of disease associations for GlycA based on complete EHRs from hospital discharge and death registries. We did not include records from primary care health centres in the analyses as the data were incompletely available and the validation of the diagnoses may vary considerably. In contrast, the validity of the nation-wide Finnish Hospital Discharge Register Diagnoses has been examined in numerous studies and found to be robust^29^. Consistent with this, we performed sensitivity analyses using predefined endpoints of major diseases defined by adjudicated compilation of hospital and reimbursement registries, and the hazard ratios were in agreement with our results using the EHR mining of hospital and death registries.

CRP is the most common marker of chronic inflammation. We therefore we aimed to evaluate if GlycA would provide additional information over and above CRP for prediction of disease onset and prognosis. GlycA associations across a majority of endpoints remained essentially unchanged when adjusted for CRP, thus demonstrating added predictive value of GlycA. Since CRP is well known predictor of morbidity and mortality in secondary prevention of cardiovascular disease^30^, we assessed whether GlycA could provide independent information on top of CRP to discriminate prognosis among patients undergoing angiography. The difference in absolute cardiovascular mortality risk was 16% between top and bottom quintile of GlycA, raising the prospect for personalized treatment strategies. Although GlycA was comparable to CRP in risk stratification in angiography patients, both biomarkers remained predictive in the multivariable model. However, further studies are required to establish whether GlycA could be used in secondary prevention in concert with CRP to assess mortality risk for angiography patients.

The question remains as to how to reduce risk for individuals with elevated GlycA. The association between GlycA, increased inflammation, and disease severity in patients with chronic disease^16–18^ in conjunction with the involvement of chronic inflammation in disease pathogenesis^23,24^ suggests the underpinnings of the associated disease risk likely lie in the low-grade chronic inflammation observed in asymptomatic adults with high GlycA levels^13^. It could be considered that GlycA is a marker of general frailty through increased inflammatory burden, and additional disease challenge posed on top of the inflammatory burden leads to poor prognosis. The recent success of the anti-inflammatory drug Canakinumab in reducing cardiovascular disease events with the unexpected benefit of reduced rates of lung cancer^21,22^ suggest anti-inflammatory treatment may be beneficial for reducing rates of other diseases predicted by elevated GlycA. Supporting this, anti-tumor necrosis factor therapy (an anti-inflammatory) was found to reduce GlycA levels, disease severity, and vascular inflammation in 16 psoriasis patients^18^. However, challenges remain in the development of anti-inflammatories for treating chronic disease, as inflammation plays a critical role in immune response and tissue repair^20^. In the Canakinumab trial increased rates of fatal infections and sepsis were observed in the treatment group^22^. Further work is required to determine strategies to mitigate the increased risk of severe infections when using Canakinumab.

Increased physical activity and weight loss are effective ways to reduce the inflammatory burden, and such lifestyle changes have also been shown to lower GlycA^31,32^. However, individuals with high disease risk profiles often do not comply with weight loss or physical activity intervention, and it is unclear whether reduction of GlycA through this approach would translate to a decrease in disease risk. Probiotics are also emerging as another approach for reducing chronic inflammation^33^, and have successfully reduced the risk for sepsis in newborns^34^. An inverse association between GlycA levels and gut microbiome species diversity has been observed in overweight pregnant women^35^, as well as with increased abundance of *Clostridales* in the gut microbiome in middle-aged men^36^.

In recent years, EHRs have emerged as a powerful tool for identification of robust biomarkers from high throughput ‘omics’ platforms in population-based cohorts. However, the underlying biology and the biomarker involvement in different diseases are often missed. This study demonstrates the validity of such an approach through the GlycA biomarker and may serve as a framework for future systems-level analyses of biomolecular data with linked EHRs in large scale cohorts such as national biobanks for a better understanding of a novel biomarker.

## Materials and Methods

### Study Cohorts

The 1997 collection of the National FINRISK Study (FINRISK97) cohort contains 8,444 individuals who responded of 10,000 randomly recruited from the five major regional and metropolitan areas in Finland to monitor the health of the adult population (aged 25–74)^25,26^. The survey included a questionnaire and a clinical examination, at which a blood sample was drawn, with linkage to national registers of cardiovascular and other health outcomes. High-throughput NMR profiling was conducted for 7,602 participants with adequate serum sample available^37^

The 2007 collection of the DIetary, Lifestyle, and Genetic determinants of Obesity and Metabolic syndrome study (DILGOM) cohort is an extension of the 2007 collection of FINRISK. In DILGOM, a detailed follow-up of 5,024 individuals was conducted to collect blood samples for omic profiling, physiological measurements, and detailed surveys of lifestyle, psycho-social, and clinical questions to study the factors leading to obesity and metabolic syndrome^27^. High-throughput NMR profiling was conducted for 4,816 participants with serum sample available. Importantly, each collection of the FINRISK study is independent, so here the DILGOM and FINRISK97 cohorts comprise independent cohorts.

The ANgiography and GEnes Study (ANGES) consists of 1000 Finnish individuals (637 men and 354 women; mean age 62)^28,38^. The patients were consequetively referred to coronary angiography because of clinically suspected coronary artery disease. High-throughput NMR profiling was conducted for 984 patients with adequate serum sample available. This study included 900 individuals with non-missing covariate data for cardiovascular mortality.

All cohort participants provided written informed consent. Protocols were designed and performed according to the principles of the Helsinki Declaration. Data protection, anonymity, and confidentiality have been assured. Ethics for DILGOM and FINRISK97 were approved by the Coordinating Ethical Committee of the Helsinki and Uusimaa Hospital District. Ethics for ANGES was approved by the local Ethics Committee of the Tampere University Hospital.

### Data quantification, processing, and quality control

Venous blood samples were collected from participants in all three cohorts. For DILGOM and ANGES venous blood was drawn after an overnight fast. For FINRISK97 the median fasting time was five hours. Serum samples were subsequently aliquoted and stored at –70°C.

GlycA levels were quantified by NMR metabolomics along with the concentrations of citrate, albumin, VLDL particle size, and 224 other measurements from serum samples for 4,816 DILGOM participants, 7,602 FINRISK97 participants, and 984 ANGES patients. Experimental protocols including sample preparation and spectroscopy are described in ref. 39. NMR experimentation and metabolite quantification of serum samples were processed by the Nightingale platform (Nightingale Health Ltd; https://nightingalehealth.com/) using a Bruker AVANCE III 500 MHz ^1^H-NMR spectrometer and proprietary biomarker quantification libraries (version 2016).

Concentrations of high-sensitivity C-reactive protein were quantified from serum samples for 4,816 DILGOM07 participants and 7,599 FINRISK97 participants using a latex turbidimetric immunoassay kit with an automated analyser.

### Electronic health records

EHR tracking was enabled by uniform diagnosis data obtained from the Finnish National Care Register for Health Care and the National Causes-of-Death Register. These registers cover all disease events, including main and side diagnoses, that have led to overnight hospitalization, or death in Finland. The EHRs are linked to study participants using their social security number, which is assigned to every permanent resident of Finland and remains the same throughout their life. EHRs were encoded according to the International Classification of Diseases 10th Revision (ICD-10) from 1996 onwards across Finland. Disease events occurring from 1987 to 1995 were encoded in ICD-9 format; these diagnoses were converted to ICD-10 format by the scheme provided by the United States Center for Disease Control Diagnosis Code Set General Equivalence Mappings (ftp://ftp.cdc.gov/pub/Health_Statistics/NCHS/Publications/ICD10CM/2011/), including combination codes. All diagnosis conversions were further verified according to the mapping scheme provided by the New Zealand Ministry of Health, National Data Policy Group (http://www.health.govt.nz/system/files/documents/pages/masterf4.xls). Manual curation of the conversion was conducted for diagnoses with mismatch in the conversion to the degree of three digits.

In this study, hospitalization and mortality diagnoses were aggregated into distinct disease “outcomes” for each individual. Each outcome corresponds to an ICD10 code at three-digit accuracy or ICD10 disease category. Main and side causes of diagnosis were treated equally in the accounting of the disease history and follow-up for each individual. EHRs were obtained for a follow-up period of 8 years for both the DILGOM and FINRISK97 cohorts to match the maximum follow-up period available for the DILGOM cohort (collected in 2007, followed prospectively until December 2015). Only the first event was considered for each outcome for each person in the prospective follow-up period. To exclude the effect of prevalent disease on associations EHRs were also obtained and aggregated as above from 1987 to sample collection of both DILGOM (events up to 20 years prior) and FINRISK97 (events up to 10-years prior); the maximum retrospective period available for EHR linkage.

To ascertain the validity of the EHR analyses, we further assessed the association of GlycA with endpoints for major common diseases (cardiovascular disease subtypes, type 2 diabetes, chronic obstructive pulmonary disease, kidney failure and any cancer) which have been carefully constructed and previously used for the FINRISK cohorts for biomarker studies^37,40^. These predefined disease endpoints were collated based on diagnoses and reimbursement information from National Health Registers: the Care Register for Health Care, the Causes-of-Death Register, the Drug Reimbursement Register, the Register for Prescribed Drug Purchases, and the Cancer Register.

To facilitate future meta-analyses, hazard ratios and standard errors are provided for all outcomes analysed in the EHR analyses in **Table S7**.

### Statistical analyses

The disease risk association analyses in DILGOM and FINRISK97 were conducted using Cox proportional hazards analyses to account for the time to first event of each diagnosis outcome. Cox proportional hazards models were fit for each outcome separately with age as time scale and adjusting for smoking, BMI, sex, systolic blood pressure, alcohol consumption as well as three all-cause mortality biomarkers previously identified alongside GlycA in FINRISK97: citrate, VLDL particle size, and albumin^6^. GlycA, albumin, citrate, BMI, systolic blood pressure, alcohol consumption and the diameter of VLDL particles were log transformed, and standardized (SD = 1) in the statistical analyses whereas current smoking and sex were used as categorical covariates. The follow-up time was matched for FINRISK97 and DILGOM to 8 years for comparability of the associations. Pregnant women were excluded from the analyses (N=100). In total 7,321 individuals from the FINRISK97 cohort and 4,540 from the DILGOM cohort were analyses in this study. The validity of Cox proportional hazards assumption was tested and all models met the assumptions.

For the incident case analysis (**Figure 1**, **Figure S1**, **Table S2**), individuals with any recorded events of the respective outcome prior to 1987 (the 10-years prior to FINRISK97 sample collection and 20 years prior to DILGOM sample collection), collectively denoted prevalent cases, were excluded from the analysis. Analysis was performed for a total of 468 disease outcomes (**Table S1**) where the number of incident cases was greater than 10 in both DILGOM and FINRISK97 in the subset of individuals who were free of prevalent cases. Meta-analysis was performed using the *metafor* R package^41^ using the inverse variance weighted fixed-effects method. The significance threshold was set to P < 1.2×10^-4^ in the meta-analysis to Bonferroni adjust for the 468 tested outcomes, with the additional requirement that the outcome was also nominally significant (P < 0.05) in both DILGOM and FINRISK97.

Sensitivity analysis to prevalent cases (**Figure S2**, **Figure S3**, **Table S3**) was performed by including prevalent cases and adding them as a covariate into the Cox proportional hazard models for each respective outcome. The minimum number of incident cases was set to 20 for the analysis including prevalent cases, as the statistical models turned out to be unstable when there were a small number of incident cases and proportionally larger number of prevalent cases. The number of disease outcomes satisfying these criteria was 376 in DILGOM, 437 in FINRISK97, and 356 in both DILGOM and FINRISK97 for the meta-analysis.

Sensitivity analysis to CRP (**Figure S4**) was performed by comparing hazard ratios from the incident case analysis to results calculated when adding CRP as a covariate to the models. CRP was log transformed and standardised in the cox proportional hazard models. Hazard ratios for the incident case analysis were re-calculated in the subset of individuals with CRP measurements for this analysis (N=4,523 in DILGOM, and N=6,985 in FINRISK97).

All-cause mortality and cardiovascular mortality (**Figure 2**) analyses in the ANGES cohort were performed using Cox proportional hazards analysis. Statistically significant covariates for all-cause mortality were age, sex, albumin, VLDL-diameter, citrate, thrombocyte count, haemoglobin levels, left ventricular hypertrophy and ejection fraction. We further tested for other risk factors which did not show association with all-cause mortality, including serum cholesterol, hyperlipidaemia, alcohol consumption, HDL cholesterol, stenosis degree over 50%, type 2 diabetes, BMI, current smoking, diastolic or systolic blood pressure. Statistically significant covariates for cardiovascular mortality were age, sex, albumin, VLDL-diameter, citrate, thrombocyte count and ejection fraction. Again, including other risk factors into the statistical model did not show association with CVD mortality risk such as serum cholesterol, alcohol consumption (non-user/moderate/high), hyperlipidaemia, HDL cholesterol, if amount of stenosis was over 50%, left ventricular hypertrophy, haemoglobin levels, type 2 diabetes, BMI, current smoking, diastolic or systolic blood pressure.

We further compared the predictive ability of GlycA and CRP for cardiovascular mortality in the ANGES cohort in separate models for the two inflammatory biomarkers, and for both included in the same model. To enable direct comparison, we limited the analyses to those individuals who had complete data on GlycA, CRP and covariates, resulting in a subsample of 582 individuals including 99 cardiovascular deaths occurring during 12-year follow-up.

Analyses were performed with R software version 3.4.1 (R Foundation for Statistical Computing; http://www.r-project.org/).

## Acknowledgements

This study was supported by funding from National Health and Medical Research Council (NHMRC) grant APP1062227 and by the Victorian Government’s Operational Infrastructure Support (OIS) program. M.I. was supported by an NHMRC and Australian Heart Foundation Career Development Fellowship (no. 1061435). JK and PW were funded by Academy of Finland (grant numbers 297338 and 307247, 312476, and 312477) and Novo Nordisk Foundation (NNF17OC0026062 and 15998). The Angiography and Genes Study (ANGES) has been financially supported by the Competitive Research Funding of the Tampere University Hospital (Grant 9M048 and 9N035), Academy of Finland: grants 286284 (for T.L), the Finnish Cultural Foundation, the Finnish Foundation for Cardiovascular Research, the Emil Aaltonen Foundation, Finland, the Tampere Tuberculosis Foundation, and EU Horizon 2020 (grant 755320 for TAXINOMISIS). V.S. was supported by the Finnish Foundation for Cardiovascular Research. M.A.-K. was supported by the Sigrid Juselius Foundation. M.A.-K. works in a unit that is supported by the University of Bristol and UK Medical Research Council (MC_UU_12013/1).

## Disclosures

PW is employee and shareholder of Nightingale Health Ltd, a company offering NMR-based metabolite profiling, including GlycA measurements. JK reports owning stock options for Nightingale Health. No other authors reported disclosures.

## Figure Legends

**Figure 1. Associations between GlycA and 8-year disease risk assessed by electronic health records**. Hazard ratios (diamonds) and 95% confidence intervals for the first diagnosis occurrence (hospitalisation or mortality) conferred per standard deviation increase of GlycA in meta-analysis of the FINRISK97 and DILGOM cohorts. The figure shows all ICD-10 categories and codes where the association with GlycA in meta-analysis was significant after multiple testing correction (P < 1.1×10^-4^; adjusting for the 468 tested outcomes) and nominally significant (P < 0.05) in both cohorts. HRs and 95% CIs are shown on a natural logarithm scale. Individual cohort results are presented in **Figure S1** and hazard ratios are detailed in **Table S2**. The spectrum of disease risk was analysed for 468 diagnosis outcomes with > 10 events in both cohorts over an 8-year follow-up. Alphanumeric codes in the square brackets indicate the ICD10 codes/categories for each outcome. Cox models were fit using age as the time scale and adjusting for sex, smoking status, BMI, blood pressure, alcohol consumption and previously discovered biomarkers for 5-year all-cause mortality (citrate, albumin, and VLDL particle size) to elucidate biology behind GlycA specific mortality risk. All available prevalent cases within prior to GlycA measurement were excluded for each association test (Methods).

**Figure 2. 12-year cardiovascular mortality associations by quintiles of GlycA in the ANGES cohort of angiography patients.** Panel A) Hazard ratios for cardiovascular mortality, relative to individuals in the lowest GlycA quintile. Cox proportional hazard models were adjusted with significant covariates from all possible tested: age, sex, albumin, VLDL-diameter, citrate, thrombocyte count and ejection fraction. Q1 indicates the lowest 20% of individuals by GlycA level, Q2 20-40%, Q3 40-60%, Q4 60-80% and Q5 >80%. Panel B) Kaplan-Meier plot of GlycA adjusted for above covariates. The subsequent residuals were used to identify top and bottom quintile of adjusted GlycA. The maximum follow-up time was 12.8 years.

